# *In silico* discovery of nanobody binders to a G-protein coupled receptor using AlphaFold-Multimer

**DOI:** 10.1101/2025.03.05.640882

**Authors:** Edward P. Harvey, Jeffrey S. Smith, Joseph D. Hurley, Alyana Granados, Ernst W. Schmid, Jason G. Liang-Lin, Steffanie Paul, Emily M. Meara, Matthew P. Ferguson, Victor G. Calvillo-Miranda, Debora S. Marks, Johannes C. Walter, Andrew C. Kruse, Katherine J. Susa

## Abstract

Antibodies are central mediators of the adaptive immune response, and they are powerful research tools and therapeutics. Antibody discovery requires substantial experimental effort, such as immunization campaigns or *in vitro* library screening. Predicting antibody-antigen binding *a priori* remains challenging. However, recent machine learning methods raise the possibility of *in silico* antibody discovery, bypassing or reducing initial experimental bottlenecks. Here, we report a virtual screen using AlphaFold-Multimer (AF-M) that prospectively identified nanobody binders to MRGPRX2, a G protein-coupled receptor (GPCR) and therapeutic target for the treatment of pseudoallergic inflammation and itch. Using previously reported nanobody-GPCR structures, we identified a set of AF-M outputs that effectively discriminate between interacting and non-interacting nanobody-GPCR pairs. We used these outputs to perform a prospective *in silico* screen, identified nanobodies that bind MRGPRX2 with high affinity, and confirmed activity in signaling and functional cellular assays. Our results provide a proof of concept for fully computational antibody discovery pipelines that can circumvent laboratory experiments.

## INTRODUCTION

Antibodies are proteins produced by B cells that recognize foreign antigens to protect against infection and are increasingly valuable as research tools and therapeutics^1^. Antibodies now make up the fastest growing class of drugs, with more than 100 antibodies approved as drugs and more than 800 in clinical trials as of 2020^2^. Antibodies can selectively differentiate between proteins that are highly similar in structure, and they can be engineered to have slow off-rates and favorable pharmacokinetic properties, permitting infrequent drug dosing^3^. Nanobodies are single-domain, heavy-chain only camelid antibody fragments and are particularly versatile due to their size, simplicity, and biochemical tractability. Nanobodies are an emerging class of antibody drugs, with one FDA-approved nanobody^4^ and others in clinical development^5^.

Most FDA-approved monoclonal antibody therapeutics were discovered using immunization technology developed in the 1970s^6^. However, many putative drug targets possess a high degree of sequence conservation in mammals, leading to immunization failures. To overcome this challenge, yeast surface display and phage display libraries provide methods to discover new synthetic antibody binders. However, these methods require specialized equipment and techniques and often generate polyreactive binders due to the lack of *in vivo* immune filtering^7^.

The recent development of AlphaFold2 and its successors has revolutionized computational and structural biology by enabling accurate predictions of protein structures and multi-protein complexes from sequence inputs^8–12^. AlphaFold2 is trained on both structural data in the Protein Databank (PDB) and Multiple Sequence Alignments (MSAs) that contain co-evolutionary information about physically interacting amino acid residues. Despite the lack of co-evolution between antibodies and antigens, we sought to investigate the possibility that AlphaFold-Multimer (AF-M), a related method trained specifically on protein complexes, can predict antibody binding to protein targets based solely on structural data in the PDB. We focused on nanobodies, which we reasoned would be ideal candidates for computational screening due to their single-chain structure that lacks a light chain region and the large number of nanobody-target complexes deposited in the PDB.

Computational antibody discovery methods raise the exciting possibility of identifying selective antibody binders completely *in silico*, circumventing challenges associated with immunization and library display selections. Computational methods like RFdiffusion have recently been reported to identify single-domain antibody fragment binders to protein targets^13,14^. However, this method performs best when structural information about similar antigen/antibody binding complexes is available and requires user-specified epitopes^13,14^.

Here, we find that AF-M reliably discriminates known GPCR-binding nanobodies from non-binding controls, even with careful efforts to separate training and test sets. Moreover, in a prospective screen of an *in silico* nanobody library of 10,000 nanobodies, we identified binders ranging in affinity from 20-200 nanomolar to MRGPRX2, a GPCR involved in non-canonical mast cell degranulation and a potential drug target for the treatment of certain inflammatory conditions. Our findings suggest that similar *in silico* antibody discovery approaches may be useful to circumvent time-consuming experimental selection campaigns and more expeditiously identify antibody binders to cellular surface receptors, particularly GPCRs.

## RESULTS

### AlphaFold-Multimer accurately predicts nanobody binding to GPCRs

AlphaFold2 and its successor AlphaFold-Multimer were trained using both sequence and structural data to generate structural predictions. By analyzing co-evolutionary patterns across homologous protein sequences in a multiple sequence alignment (MSA), residue interactions that contribute to structural stability or function can be inferred. This co-evolutionary signal, in combination with experimentally determined protein structures, has allowed AlphaFold2 and related methods to achieve near-experimental accuracy for many proteins^10–12^. However, the performance of these models is notably diminished for antibody-antigen pairs, as these pairs do not meaningfully co-evolve, and the models must rely exclusively on patterns in the available protein structure data. Despite this limitation, we observed that AlphaFold-Multimer could accurately predict the unusual structure of the GPCR angiotensin II type 1 receptor (AT1R) bound to a synthetic nanobody, AT118-H (**Figure 1A, 1B**). Importantly, this experimentally determined cryo-EM structure was deposited to the PDB after the model’s training cutoff date^15^. The strong agreement between the predicted and experimentally determined structures (RMSDs: overall = 2.9 Å; receptors = 2.9 Å; nanobodies = 1.8 Å; CDR1 = 1.52 Å, CDR2 = 2.65 Å, CDR3 = 1.54 Å) is especially surprising because AT118-H induces an unusual, previously unseen conformation of AT1R in which the external side of the receptor is in an active-like state while the internal side of the receptor is in an inactive-like state^15^. The correct prediction of this unusual state shows that AF-M infers nuanced structural features rather than merely recapitulating nanobody-GPCR structures in its training data. Furthermore, AF-M’s confidence metrics for this interaction pair are quite high, as reflected by the low AT1R/AT118-H interface predicted aligned error (PAE) values (**Supplementary Figure 1C**). In recent years, the number of structures of nanobodies bound to GPCRs has increased rapidly, growing from a handful in 2015 to more than 150 deposited structures in 2023 (**Figure 1C, Supplementary Figure 1A-B**). This might, in part, explain why AF-M correctly predicted the structure of AT1R bound to AT118-H, while also raising the possibility of continued improvements in the future.

**Figure 1.**
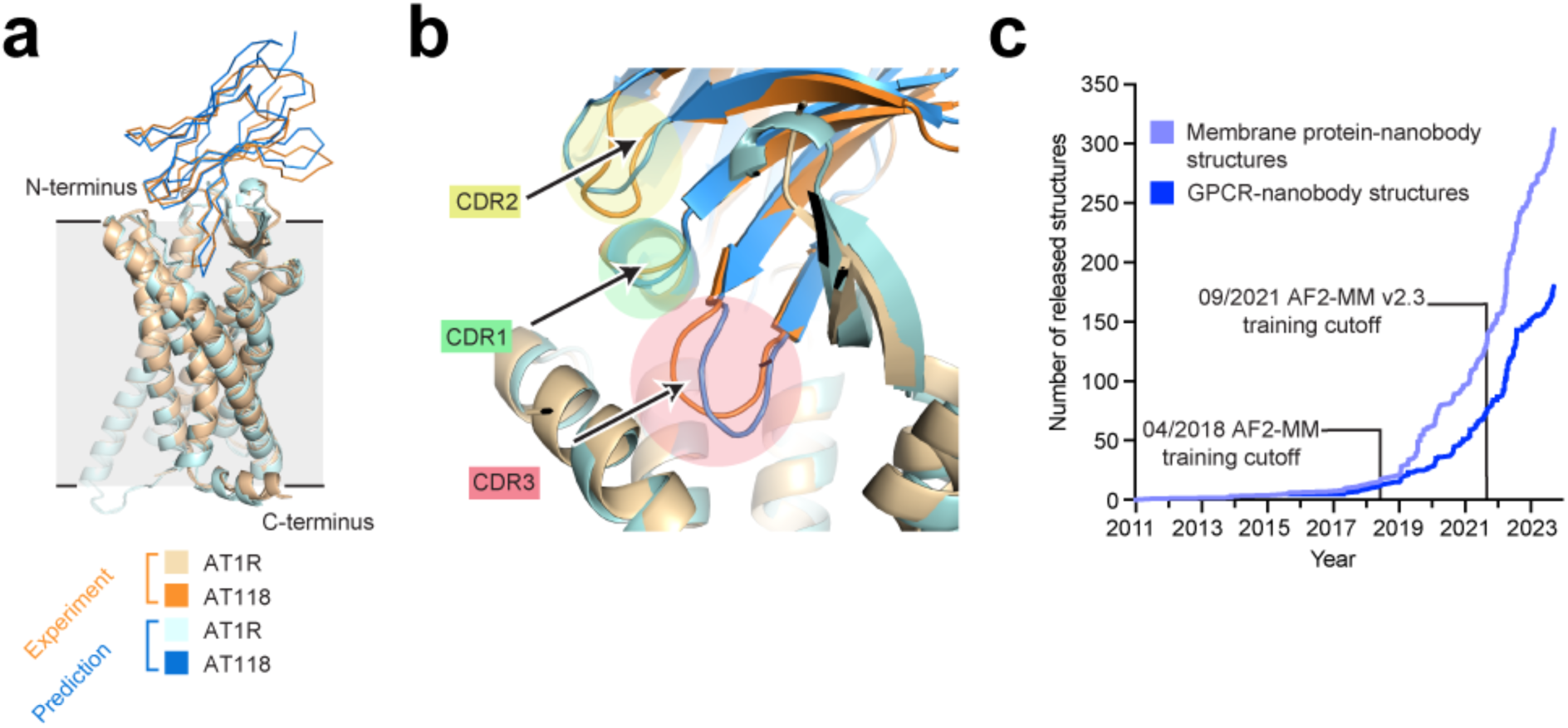
AlphaFold-Multimer accurately predicts an unusual binding pose and structure of a nanobody with a GPCR. a) Overlay of the experimentally determined structure of the complex of the GPCR AT1R with nanobody AT118-H (PDB 8TH3) and the AF-M prediction demonstrating nearly identical structural models. b) The CDR regions of AT118-H in the experimentally acquired and computationally predicted structural models are closely aligned. c) The number of released nanobody structures bound to either membrane proteins or GPCRs has increased rapidly.

The surprising ability of AF-M to accurately predict the structure of the AT1R/AT118-H complex led us to ask how generalizable this result might be. Specifically, we sought to test whether AF-M could distinguish true nanobody binders of GPCRs from non-binding controls. We also sought to assess performance on non-GPCR membrane proteins and soluble protein targets. We compiled sets of known nanobody-antigen pairs that were supported either by direct binding data or experimentally solved structures deposited to the PDB after the AF-M training cutoff date of 9-30-2021 (**Extended Data Tables 1-3**). Our training sets include 30 validated GPCR nanobody binders, 18 membrane protein nanobody binders, and 49 soluble protein nanobody binders (Extended Data Tables 1-3). To generate sets of non-binding pairs, we permuted nanobody-antibody pairs such that nanobodies were paired with antigens other than their reported antigen. To perform predictions in a high throughput manner, we utilized a modified AF-M ColabFold script optimized for faster prediction speed and decreased memory storage requirements^8,16,17^, and we generated structural predictions of these curated nanobody-antigen pairs to investigate which outputs, if any, were predictive of nanobody binding.

**Table 1:** MRGPRX2 binding characteristics for the three leading candidate nanobodies in HEK293T cells and ROSA mast cells.

|  | HEK293T cells |  | ROSA mast cells |
| --- | --- | --- | --- |
| | Kd<br>(nM ± SEM) | Bmax<br>(Relative ± SEM) | Kd<br>(nM ± SEM) |
| Nanobody Sim8619 (rank 1) | 200 ± 20 | 97% ± 3 | 100 ± 10 |
| Nanobody Sim9877 (rank 5) | 160 ± 30 | 100% ± 6 | 20 ± 4 |
| Nanobody Sim4784 (rank 7) | 80 ± 30 | 43% ± 5 | 50 ± 10 |

We then developed a parser that extracted metrics that we hypothesized might differ between interacting and non-interacting pairs. Several global metrics were calculated (e.g., average predicted template modeling score (pTM) across the five AF-M models). Additionally, we filtered for residue pairs near the nanobody-target interface (defined as intra-chain residue pairs for which the Cα atoms were ≤10 Å from each other) and used these data to calculate several interface-specific metrics. Among these were average interface predicted aligned error (PAE), average interface predicted local distance difference test (pLDDT), pDockQ^18^; and average model support^16,18^. In addition to metrics averaged across all five models, metrics derived solely from the highest confidence model (as ranked by AF-M) were also considered.

We found that several of these metrics were predictive of true binding in our GPCR-nanobody dataset (**Figure 2A)**. In particular, seven metrics had areas under the receiver operating characteristic curve (AUROCs) above 0.65: average pTM across five models (AUROC = 0.73), best model pTM (AUROC = 0.71), average interface PAE (AUROC = 0.69), best model interface PAE (AUROC = 0.68), average interface pLDDT (AUROC = 0.67), best model interface pLDDT (AUROC = 0.65), and best model pDockQ (AUROC = 0.66). We combined these features (except best model pDockQ, which is largely derived from best model pLDDT and therefore redundant) by scaling them to span the range 0– 1 and calculating the product of all six scaled features to generate a Linear Combination Feature (LCF). This combination feature had an AUROC of 0.71, which is slightly lower than that of its highest component feature, but which we reasoned may be more robust than a single feature chosen based on a modestly sized benchmarking set.

**Figure 2.**
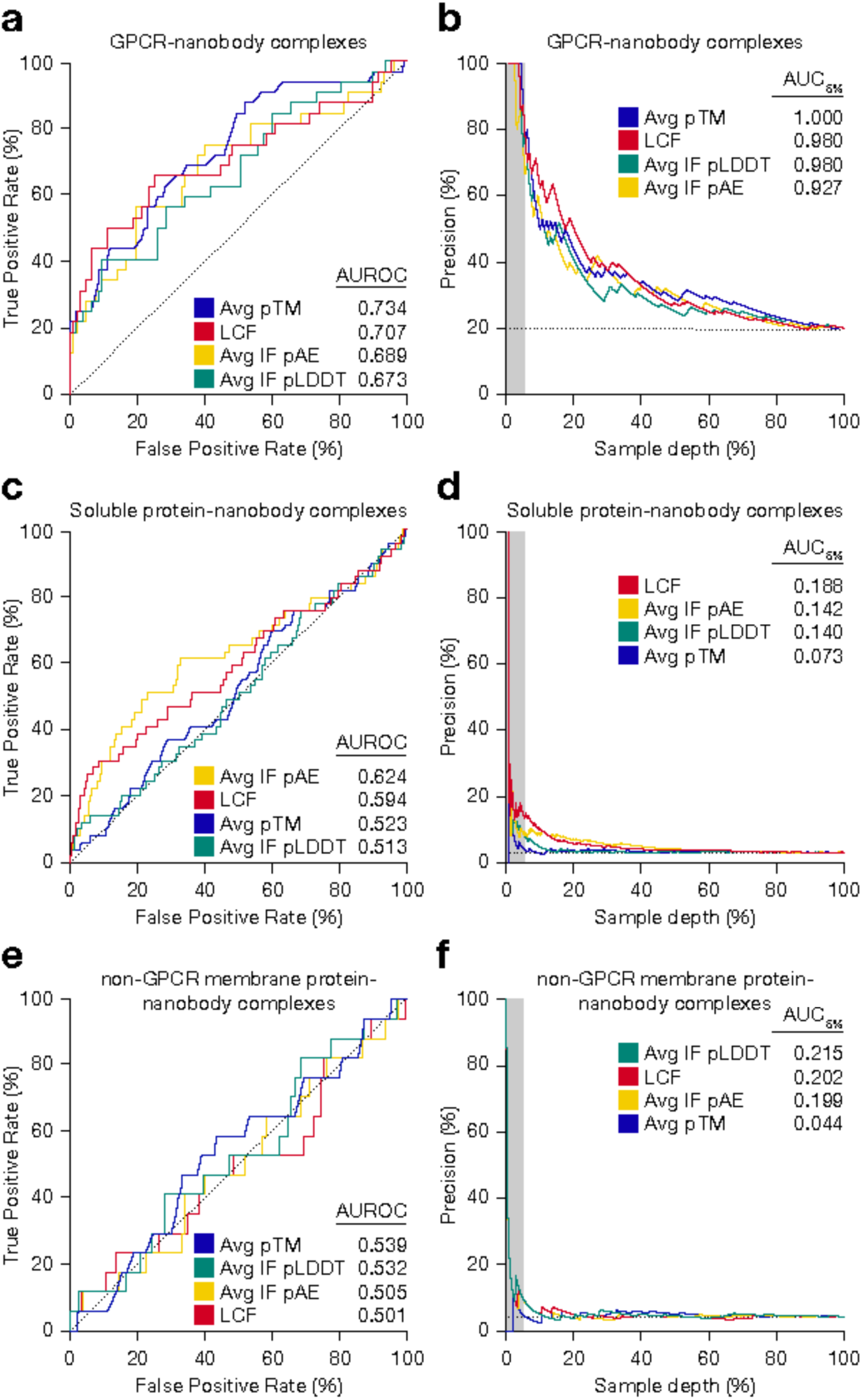
AF-M metrics differentiate between validated nanobody GPCR binders and negative controls. a) Receiver operating characteristic curves comparing the ability of select AF-M confidence metrics and the Linear Combination Feature (LCF) to differentiate between true nanobody GPCR binders and negative controls. b). Comparison of AUC5% values for AF-M confidence metrics and the LCF, illustrating performance specifically on highest ranked nanobodies. c-f) AF-M confidence metrics and the LCF possess poor ability to differentiate between true nanobody soluble protein binders and negative controls (c-d) and true nanobody non-GPCR membrane protein binders and negative controls (e-f).

In contrast to its predictive ability to differentiate true GPCR-binding nanobodies from negative controls, AF-M is currently unable to reliably differentiate positive and negative control pairs for soluble protein-nanobody interactions and non-GPCR membrane protein-nanobody interaction pairs, as illustrated by the previously described metrics having AUROCs close to 0.5 **(Figure 2C, 2E)**. Recently, AlphaFold3 was reported to possess an improved ability to predict antibody binding to proteins^9^. We used our benchmarking sets to compare the predictive ability of AlphaFold3 and AF-M run once using single initialization seeds for nanobodies and found that the AUROCs for ipTM metrics were nearly identical for the GPCR benchmarking set (**Supplementary Figure 2A**) and were moderately improved for the soluble protein benchmarking set (**Supplementary Figure 2B**). Similar to AF-M, AlphaFold3 was not currently able to differentiate between true non-GPCR membrane protein nanobody binders and negative controls (**Supplementary Figure 2C).** Finally, we assessed ESMfold, an orthogonal protein structural prediction model, and found that ESMFold is unable to differentiate between true positive binding interactions and negative control binding pairs for our GPCR benchmarking dataset (**Supplementary Figure 2D-F)**^19^. These results show that AF-M is currently more proficient at predicting nanobody binding to GPCRs than other structural prediction algorithms tested.

The ability of AF-M to accurately rank binding vs. non-binding GPCR nanobodies raised the possibility that AF-M could be used as a virtual screening tool for antibody discovery. In such a screen, presumably only highly-ranked hits would be selected for further experimental validation. Therefore, model precision (i.e., the proportion of the model’s “positive” classifications that are truly positive) for the highest ranked nanobodies is more pertinent than model accuracy for the average nanobody in the dataset, which is reflected by metrics like AUROC. To assess performance specifically on the highest ranked nanobodies, we calculated precision as a function of sample depth (the assessed proportion of the distribution of predictions ranked from highest to lowest according to their AF-M confidence metrics) and calculated AUC5% values, defined as the precision for only the highest-ranked 5% of nanobodies by a given metric **(Figure 2B, 2D, 2F)**. The linear combination feature and each of its component metrics exhibited stellar AUC5% values for the GPCR benchmarking dataset (ranging from 0.93–1). However, the AUC5% metrics for the soluble and membrane protein benchmarking sets were much lower (≤0.22). These results suggest that AF-M could currently be deployed for *in silico* discovery of nanobody binders to GPCRs, but not soluble or non-GPCR membrane proteins.

### *In silico* discovery of MRGPRX2 nanobodies

To perform a virtual nanobody screen to find MRGPRX2 binders, we generated ten thousand simulated nanobody sequences that match the design parameters of our previously published nanobody yeast display library^20^ (**Figure 3A**). As our candidate target GPCR, we selected MAS-related GPR family member X2 (MRGPRX2), a GPCR that regulates IgE-independent mast cell degranulation. This receptor was chosen because its shallow binding pocket suggests a more forgiving thermodynamic landscape relative to other GPCRs^21^. MRGPRX2 is found exclusively in primates and has low sequence similarity to MRGPRX1, MRGPRX3, and MRGPRX4, which may reduce certain contributions from other GPCRs within the PDB to the AF-M training algorithm. Currently, no FDA-approved medications target MRPGRX2, making it an attractive therapeutic target.

**Figure 3.**
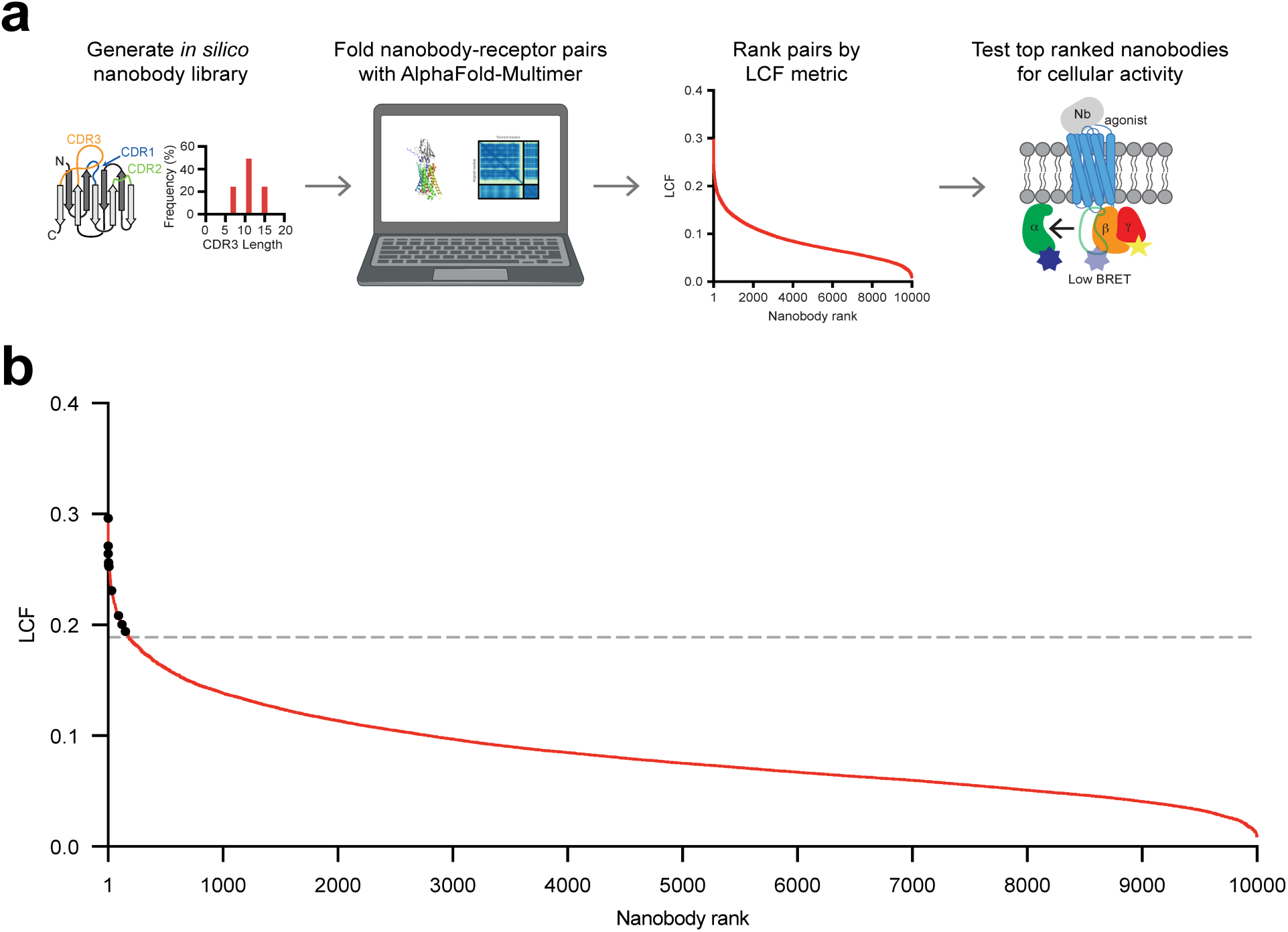
**Deploying AF-M *in silico* screening for nanobody binders to the GPCR MRGPRX2**. a) Flowchart describing the *in silico* nanobody screening process. Ten thousand nanobody sequences that met prior library design specifications (McMahon *et al.* 2018) were computationally co-folded with the GPCR MRGPRX2 in AF-M. Nanobodies were then ranked by the LCF metric. A sample of the highest ranked nanobodies were expressed, purified, and tested for binding and activity in signaling assays. b) Co-folding ten thousand nanobodies with the GPCR MRGPRX2 in AF-M generated 179 nanobody-receptor pairs with LCF values greater than that of the highest value in the non-binding GPCR-nanobody set (indicated by dashed gray line). Black circles indicate nanobodies prioritized for validation studies.

We predicted 10,000 MRGPRX2-nanobody structures using AF-M without templates to prevent AF-M from being unduly biased by a single structure and subsequently ranked the predictions according to our LCF metric. Importantly, structures of MRGPRX2 were deposited to the PDB several months after the AF-M training cutoff, so specific structural details of MRGPRX2 were not available *a priori*^22,23^. Nanobodies that possessed potential liabilities, such as glycosylation sites within the CDRs or CDR sequences predicted to be polyreactive were removed^7^. 179 nanobodies (1.79% of total simulated nanobodies) demonstrated compelling LCF metrics, i.e., an LCF greater than the highest LCF of the negative control nanobodies in the GPCR nanobody benchmarking dataset **(Figure 3B)**. The top six ranked nanobodies with no liabilities were expressed and purified as Fc fusions **(Supplementary Table 1)**. In addition, four lower-ranked nanobodies distributed evenly throughout the top 179 were chosen for expression and purification to broadly sample the population of *in silico* hits.

We first screened 10 purified candidate nanobodies for binding to an immortalized mast cell line (ROSA) that endogenously expresses MRGPRX2 at high levels, which we confirmed by flow cytometry (**Figure 4A**). Nanobody Sim8619 (rank 1), Sim9877 (rank 5), Sim4784 (rank 7), and Sim4177 (rank 90) showed high levels of binding to ROSA cells, while our negative control nanobody GPCR binder, Nb 60, showed no appreciable binding^24^ **(Figure 4B)**. Sim8619 (rank 1), Sim9877 (rank 5), and Sim4784 (rank 7) demonstrated reasonably monodispersed size exclusion chromatography profiles and were selected for further validation experiments. However, Sim4177 (rank 90) was poorly behaved biochemically **(Supplementary Figure 4A, 4B)** and was therefore excluded from future analyses. We further measured candidate nanobody binding to mast cells in dose-response format. Sim8619, Sim9877, and Sim4784 bound ROSA mast cells in a dose-dependent manner **(Figure 4C-E)**. In complementary experiments, HEK293T cells (which lack endogenous MRGPRX2) were transfected with either MRGPRX2 or empty vector. Nanobody Sim8619, Sim9877, and Sim4784 showed similar binding properties and estimated dissociation constants as ROSA mast cells, providing evidence that the nanobodies are binding specifically to MRGPRX2 **(Table 1) (Supplementary Figure 5)**.

**Figure 4.**
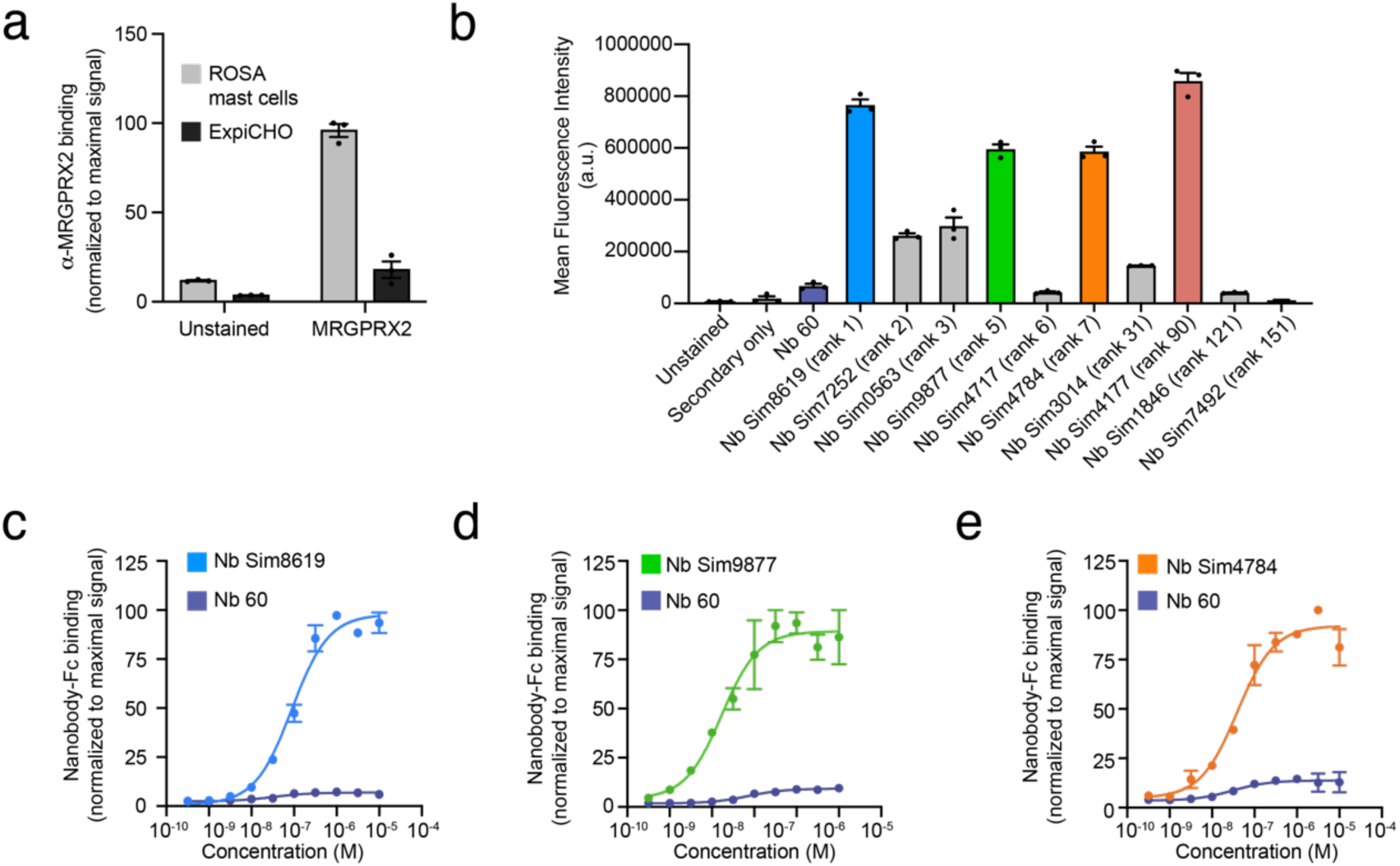
Binding of candidate MRGPRX2 nanobodies to mast cells. a) MRGPRX2 expression levels were assessed in ROSA mast cells expressing MRGPRX2 and ExpiCHO cells lacking MRGPRX2 using a commercial anti-MRGPRX2 antibody (Biolegend Cat# 359005, AB_2750139). b) Flow cytometry screen of top-ranked clones to ROSA mast cells. c) Nanobody Sim8619 (rank 1), d) nanobody Sim9877 (rank 5), e) nanobody Sim4784 (rank 7) binding to ROSA mast cells relative to Nb 60 negative control. Panels a-b; error bars represent mean +/-SEM of technical replicates performed in triplicate. Panels c-e; experiments were performed in biological duplicate with error bars representing mean +/-SEM. A representative gating strategy is shown in **Supplementary Figure 7**.

Next, we characterized the functional activity of the three putative MRGPRX2 binders. Activation of MRGPRX2 leads to mast cell degranulation^25,26^, and we used an established β-hexosaminidase release assay to measure the extent of degranulation upon the addition of the nanobodies to mast cells^27^. Known MRGPRX2 small molecule agonist Compound 48/80 induced the release of approximately 70% of granules from mast cells (**Figure 5A**). The addition of Sim8619 (rank 1), Sim9877 (rank 5), or Sim4784 (rank 7) did not result in an increase in degranulation over the vehicle or Nb60 negative control, indicating that these nanobodies are not functional agonists of MRGPRX2 **(Figure 5A)**. Next, we tested if Sim8619 (rank 1), Sim9877 (rank 5), and Sim4784 (rank 7) were functional antagonists by assessing their ability to block compound 48/80 mediated degranulation. While the negative control nanobody, Nb60, had no effect on compound 48/80 mediated degranulation, the addition of Sim8619 (rank 1), Sim9877 (rank 5), or Sim4784 (rank 7) attenuated 48/80-induced degranulation, suggesting they are functional antagonists **(Figure 5B**).

**Figure 5.**
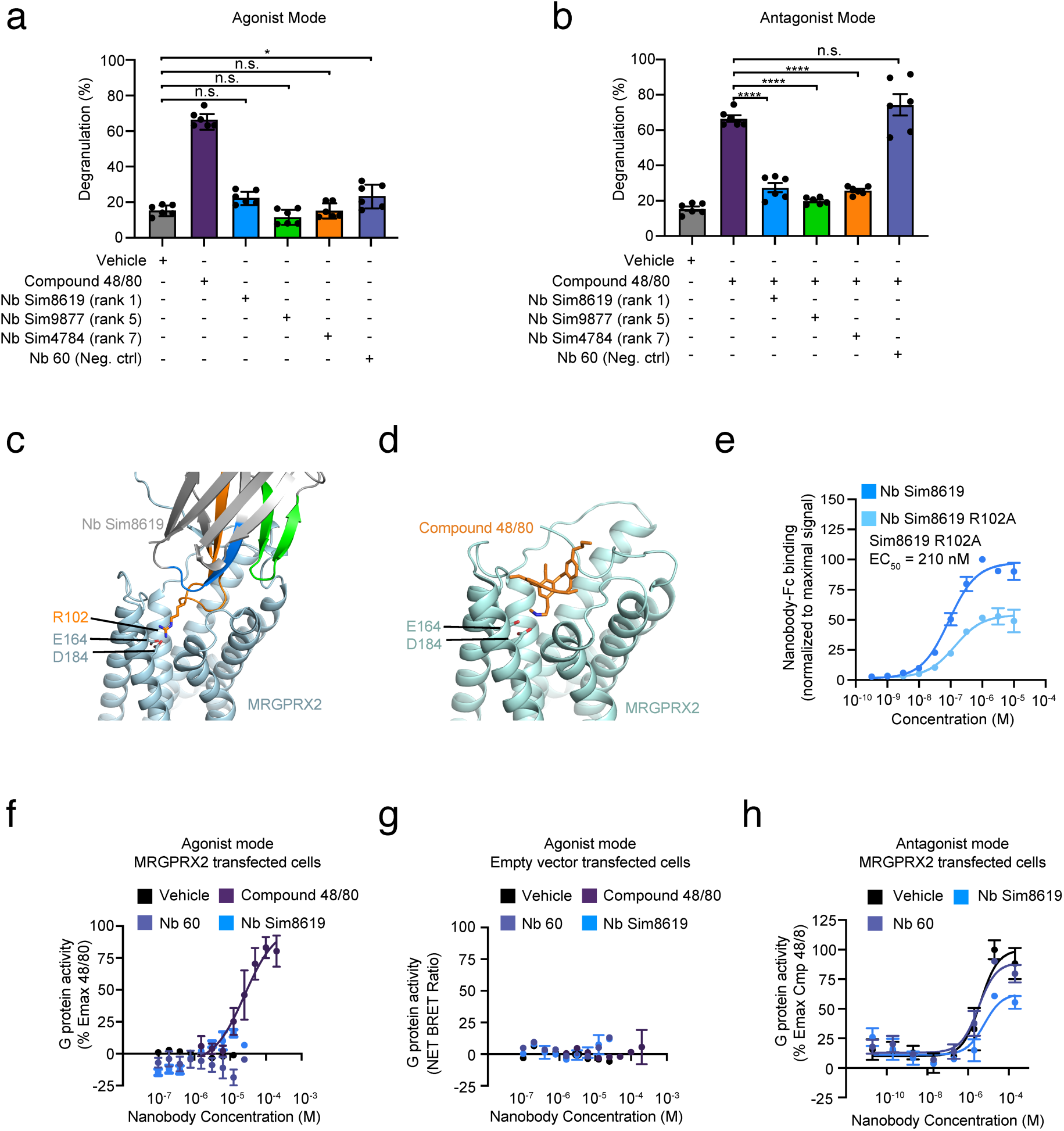
Functional characterization of MRGPRX2 nanobody binders. a) “Agonist mode” degranulation assay, where degranulation of ROSA cells from Sim8619 (rank 1), Sim9877 (rank 5), and Sim4784 (rank 7) treatment was compared to the positive control agonist 48/80 or the negative control Nb60. b) “Antagonist mode” degranulation assay, where ROSA mast cells were pre-treated for 15 min with either Sim8619 (rank 1), Sim9877 (rank 5), and Sim4784 (rank 7), then treated with compound 48/80 (50 DM). c) R102 of Sim8619 (rank 1) is predicted in the AF-M structural model to make electrostatic interactions with two acidic residues (E164 and D184) on MRGPRX2. d) The small molecule agonist compound 48/80 occupies the same binding site on MRGPRX2 as Sim8619 (rank 1) is predicted to bind, interacting with E164 and D184 (PDB 7VV6). e) Nanobody Sim8619 (rank 1) with an R102A mutation binds to ROSA mast cells with a weaker EC_50_ (210 nM) and lower maximal binding value compared to wild type Sim8619. f-g) G protein activation as assessed by TRUPATH Gi heterotrimer BRET dissociation assay in HEK293T cells overexpressing TRUPATH Gi BRET constructs and MRGPRX2 (f) or TRUPATH Gi BRET constructs and empty vector (pcDNA 3.1) (g). Sim8619 (rank 1) did not cause appreciable G protein activation relative to the positive control agonist 48/80. h) HEK293T cells overexpressing TRUPATH Gi BRET constructs and MRGPRX2 were pretreated with Sim8619 (rank 1), negative control antibody nanobody 60, or vehicle for 45 minutes, and subsequently treated with the indicated concentration of MRGPRX2 agonist 48/80. Experiments were performed in at least biological duplicate. Error bars indicate mean +/-SEM of biological replicates. Mean +/-SEM is shown in all panels. For panels A and B, six biological replicates were performed and included at least two separately prepared nanobody purifications. Error bars indicate mean +/-SEM of six biological replicates. Statistical analysis was performed using ANOVA, and a Dunnett’s post hoc test was performed comparing all samples to vehicle treatment ****p < 0.0001, *** p < 0.0002, ** p <0.0021, * p < 0.0332, n.s. not significant. For panel h, *p < 0.05 two-way ANOVA, main effect of nanobody pretreatment.

Interestingly, AF-M predicts Sim8619 (rank 1), Sim9877 (rank 5), and Sim4784 (rank 7) bind in the same orthosteric pocket site as compound 48/80 **(Supplementary Figure 6).** The CDR3s of nanobody Sim8619 (rank 1) and nanobody Sim 4784 (rank 7) mimic the binding interaction of a positively charged side group of the MRGPRX2 agonist compound 48/80 with two acidic residues, E164 and D184, on the receptor **(Figure 5C and 5D, Supplementary Figure 6)**, which have previously been shown to be essential for compound 48/80’s agonist activity^28–30^. To test this predicted binding pose, we mutated a critical arginine residue (R102A) within our top-ranked nanobody, Sim8619 (rank 1), which the AF-M predicted alignment suggested would disrupt binding with E164 and D184 in the MRGPRX2 orthosteric site **(Figure 5C, 5D).** In support of this binding pose, the R102A mutant bound to ROSA cells with a two-fold decreased affinity and lower maximal binding **(Figure 5E)**. The predicted similarity of these binding interactions provides a structural rationale for the ability of Sim8169 to block compound 48/80 mediated degranulation.

Based on these predictions and binding data, we proceeded to test if Sim8169 (rank 1) could directly compete with the MRGPRX2 agonist 48/80 in its ability to activate the G protein Gi, an established signaling output of MRGPRX2^31^. We first tested the ability of Sim8619 to independently activate Gi. Unlike compound 48/80, which robustly activated Gi in an MRGPRX2-dependent manner, Sim8619 did not, with only slight G protein activity only at high concentrations relative to the agonist compound 48/80 **(Figure 5F, FG)**. However, pretreatment of MRGPRX2-expressing cells with Sim8619 prior to 48/80 treatment lowered the G protein activity Emax and slightly right-shifted the EC_50_ **(Figure 5H)**. This competition assay provides further support for the predicted binding site of Sim8169 at MRGPRX2, as well as further support for Sim8169 acting as a functional MRGPRX2 antagonist.

## DISCUSSION

Small molecule docking is widely used to identify compounds that bind to proteins of interest. *In silico* compound docking helps to make laboratory work more efficient by enriching for compound binders compared to naively screening chemical space. Similarly, here we show that AF-M can be deployed to screen GPCR-binding nanobodies by ranking nanobody-GPCR interactions according to their AF-M confidence metrics. Here, we show that AF-M can accurately identify true nanobody binders to the GPCR MRGPRX2. We identified three nanobody binders to MRGPRX2 and validate that these nanobodies have functional antagonist activity. Despite the improved capabilities of AF-M over its predecessors, these findings were in some ways surprising, as unlike many other protein-protein interactions, co-evolution between nanobodies or antibodies and GPCRs does not occur. This lack of co-evolution between antibodies and their binding partners limits the utility of one of the core metrics used in AF-M predictions. In recent years, the number of structures of nanobodies bound to GPCRs has increased rapidly, increasing from a handful in 2015 to more than 150 deposited structures in 2023 **(Figure 1C)**. This rapid increase in reliable training data may, in part, explain our success with a GPCR and not with other protein targets, as AF-M has more examples within its training set to utilize. Future iterations of AF-M or similar models are likely to continue to improve the ability to recognize antibody binders accurately.

We chose MRGPRX2 as a test candidate because of its clinical potential and because it is a known promiscuous binder, thereby making small library screening more likely to succeed. The shallow binding pocket accommodates a broad range of charged ligands^32^. Further studies are needed to determine if this strategy can be deployed similarly for other GPCRs. In our study, nanobody Sim8619 (rank 1), nanobody Sim9877 (rank 5), and nanobody Sim4784 (rank 7) are nanomolar binders and function as MRGPRX2 antagonists, as assessed by blocking mast cell degranulation. Derivatives of these nanobodies could be of therapeutic interest for diseases such as chronic spontaneous urticaria^33^.

More broadly, antibodies have emerged as essential reagents for deciphering GPCR biology. Approximately 35% of FDA-approved drugs target GPCRs, highlighting that pharmacologic modulation of GPCRs is a critical cornerstone of modern medicine^34^. Antibodies can offer certain advantages over small molecules targeting GPCRs in many cases, such as extended half-life and often increased specificity. However, as of 2021, few GPCR-targeting antibodies are currently in clinical development, with only three GPCR-targeting monoclonal antibodies approved and less than 1% of the GPCR therapeutic pipeline consisting of antibodies^35^. This deficit is in large part due to the technical challenges associated with conducting GPCR antibody discovery campaigns. GPCRs are conformationally dynamic and often poorly expressed, making it particularly challenging to purify the quantities of biochemically stable and homogenous samples needed for traditional antibody discovery campaigns.

For more challenging *in silico* targets, Bayesian optimization methods such as active learning can be deployed to learn the binding contributions of specific amino acids from a subset of antibodies screened to guide the design of remaining library members and more efficiently explore antibody chemical space^36^. As more structures of antibodies bound to proteins continue to be deposited to the PDB, the predictive ability of AF-M and other computational models, such as RFdiffusion, will likely increase, further improving the already noteworthy utility of this method to discover new antibody fragment binders and decipher other protein-protein interactions.

### Code Availability Statement

Code for analyzing AF-M results and ranking nanobody clones can be found on GitHub: https://github.com/kruselab/MRGPRX2-AF-M-screen.

## STATISTICAL METHODS

Prism software (Graphpad) was used to analyze data and perform error calculations. Data are expressed as arithmetic mean ± SEM.

## REPORTING SUMMARY

Further information on research design is available in the Nature Portfolio Reporting Summary linked to this article.

## ACKNOWLEDGEMENTS

This work was funded by a Christopher Walsh Postdoctoral Fellowship to E.P.H.; funding and support provided by the National Institutes of Health Dermatology Training grant 5T32AR007098, Arthritis and Musculoskeletal and Skin Diseases K08AR084617, and the Dermatology Foundation to J.S.S.; NIH training grant 5T32GM007226-46 to J.D.H; support from the National Science Foundation (DGE 2140743) to E.W.S.; NIH grant HL098316 to J.C.W.; J.C.W. is an American Cancer Society Research Professor and a member of the Howard Hughes Medical Institute; NIH grant DP5OD036136, a UCSF Sandler Fellowship, and a Program in Breakthrough Biomedical Research award to K.J.S.; NIH TR01 grant R01CA260415 to A.C.K. We thank Meredith Skiba for helpful discussions and experimental support, Justin English for TRUPATH integrated plasmids, Tracy Lou for technical assistance, and Edward Harvey Jr., Louis Hollingsworth, Daniel Same-Guerra, and James Osei-Owusu for assistance with AlphaFold3 *in silico* screening. We thank Xiaojing Cong (Institut de Génomique Fonctionelle) for helpful discussion about AlphaFold prediction for nanobody-GPCR complexes. We thank Daniel Gray for helpful discussions regarding simulation library generation.

## AUTHOR CONTRIBUTIONS

E.P.H., K.J.S., J.S.S., and A.C.K. designed experiments. E.P.H., K.J.S., J.S.S., and A.C.K. wrote the manuscript with input from all authors. E.P.H., K.J.S, J.S.S., A.G., J.L.-L., E.M.M., and M.P.F. performed biochemical experiments. V.G.C-M. provided experimental support and helpful discussions. E.P.H. ran AlphaFold simulations. J.D.H. and E.W.S. wrote code to analyze AF-M outputs. E.P.H., K.J.S., J.S.S., J.D.H., S.P., and A.C.K. analyzed data. S.P. conducted ESMFold simulations. D.S.M., J.C.W., K.J.S., and A.C.K. supervised the work and provided financial funding.

## Methods

### AF-M and AlphaFold3 multimer predictions

AF-M structural predictions were generated using a modified local ColabFold script (https://github.com/YoshitakaMo/localcolabfold) operated on a Lamba Labs server with a NVIDIA A100 GPU^8,10^. In particular, AF-M multimer v2 or v3 was used with three recycles and was run without templates and amber processing^12^. PyMol was used to visualize AF-M multimer structural predictions, and confidence in predictions was assessed by contact analysis scripts. AlphaFold3 structural predictions were generated using the AlphaFold3 server (http://alphafoldserver.com) with standard settings, and ipTM/pTM values were obtained from the server output^9^.

### ESMFold multimer predictions

ESMFold binding complex structure predictions were generated using standard run scripts with the multimer setting (https://github.com/facebookresearch/esm/tree/main#esmfold)^19^. Models were run using a single NVIDA TeslaV100s GPU. Predicted structures contained predicted-LDDTs (pLDDTS) which were parsed from the output.pdb file. Average pLDDTs of the residues in the full VHH, all the CDR regions, and just CDR3 were used as binding scores from ESMFold. To retrieve the CDR annotations, IMGT numbering of the nanobody sequences in the folding benchmark was performed using ANARCI^37^.

### Recombinant nanobody expression and purification

To recombinantly express nanobodies fused to human IgG1 Fc, nanobody DNA sequences followed by a (GGS)3 linker were cloned into pFUSE-CHIg-hG1 (InVivo Gen) containing a H435A substitution, which prevents IgG Fc binding to Protein A resin ^38,39^.

Expi293F cells were then transiently transfected with nanobody plasmids. Briefly, 200 mL of Expi293F cells cultured in Expi293 expression media (Thermo-Fisher) were grown to a density of 3×10^6^ cells/mL. Then, cells were transiently transfected using nanobody DNA (0.16 mg total DNA) and FectoPro transfection reagent (Polyplus) at a 1:1 DNA/FectoPro ratio. 16 hours after transfection, cells were enhanced with 3 mM Valproic acid sodium salt (Sigma-Aldrich) and 0.8% D-(+)-Glucose (Sigma-Aldrich). Transfected cells were cultured for 6 days to produce nanobody Fc-fusions. Then, the media was separated from cells by centrifugation at 4000xg for 15 minutes at 4 °C, and applied to protein A resin equilibrated with 20 mM HEPES, 150 mM NaCl (pH 7.5). The protein A resin was then washed with 20 column volumes 20 mM HEPES, 150 mM NaCl (pH 7.5). Following resin washing, nanobody IgG1 Fc fusions were eluted from the resin using 100 mM citrate (pH 3) directly into 2 M HEPES (pH 8) and the pH of the eluted and neutralized protein solution was checked using pH strips. Eluted nanobody Fc fusions were then dialyzed overnight in 20 mM HEPES, 150 mM NaCl, 10% glycerol (pH 7.5) and were flash frozen. Protein purity was assessed by SDS-PAGE and analytical size exclusion chromatography runs on a Superdex 200 Increase 3.2/300 gel filtration column (GE Healthcare).

### G protein signaling assays

The Gi TRUPATH bioluminescence energy transfer (BRET) integrated plasmid was a gift from Justin English. Assays were conducted similar to those previously described^40–42^. Briefly, HEK293T cells were transfected with a human N-terminal FLAG-tagged MRGPRX2 in a pcDNA3.1 expression vector or empty vector control and a Gi TRUPATH plasmid. After 24 hours, cells were plated in a 96-well plate in phenol red free DMEM supplemented with 2% FBS, 1% Glutamax, and 1% antibiotic-antimycotic (Sigma).

Approximately 48 hours after transfection, media was replaced with assay buffer (HBSS without calcium or magnesium + 20 mM HEPES + 3 µM coelenterazine-400a (Nanolight) and BRET ratios were obtained using a Promega plate reader. For agonist mode, data were normalized to a pre-read prior to nanobody treatment. For the initial MRGPRX2 activation screen, cells were treated with 20 µM of each MRGPRX2 simulation nanobody and read approximately 10 minutes after addition. For competition assay (“antagonist mode”), cells were pretreated with 10 µM of the indicated nanobody or vehicle for 45 minutes, and then a pre-read was conducted. Cells were then treated with variable concentrations of the MRGPRX2 agonist compound 48/80 or vehicle, and the net BRET ratio was calculated by first subtracting the pre-read from the post-read, then subtracting vehicle pre-treatment, vehicle treatment condition from other treatment conditions. Independent experiments were conducted in technical triplicate and merged for a biological replicate, with the indicated number of biological replicates per experiment included within the pertinent figure legends.

### Nanobody on-cell cytometry binding assays

The wild type ROSA mast cell line^43^ was a gift from the Galli lab (Stanford University) and was confirmed negative for mycoplasma. Rosa cells were cultured at 37 °C and 5% CO_2_ in IMDM supplemented with 1% Penicillin-Streptomycin (Pen-Strep), 1% sodium pyruvate (GIBCO), 1% MEM (minimal essential medium) (GIBCO), 2% MEM non-essential amino acids (GIBCO), 1% L-glutamine (GIBCO), 1% insulin transferrin-sodium selenite (GIBCO), 0.3% bovine serum albumin, with 80 ng/mL fresh mouse stem cell factor (SCF) (R&D Systems). To obtain nanobody dose response curves, cells were harvested by centrifugation at 1,000xg and washed twice with cold phosphate buffered saline (PBS).

The cells were then counted, and 100,000 cells were added to each well of a V-bottom 96 well plate (Grenier Bio). Cells were washed again with 100 μL cold PBS and then were incubated with different concentrations of either the candidate nanobodies, commercial MRGPRX2 antibody (Biolegend Cat# 359005, AB_2750139), or the negative control Nb 60-Fc fusion (β2AR internal nanobody binder negative control), or buffer alone for thirty minutes at 4°C^24^. Following another PBS wash, cells were incubated with 1 μg/mL anti-human IgG Fc-488 (Invitrogen Cat#A-11013) for 30 minutes at 4°C. After secondary antibody incubation, cells were washed twice more with PBS and were analyzed by flow cytometry using a CytoFLEX cytometer (Beckman-Coulter). In detail, live/dead gates were set using SSC-A versus FSC-A and singlet gates using FSC-H versus FSC-A, and fluorescence intensity at 488 nm was recorded in the doubly gated population. Mean fluorescence intensity (MFI) at 488 nm was plotted in GraphPad Prism as an assessment of nanobody binding to cells. For Kd calculations, non-specific signal was MFI in the nanobody 60 condition.

For HEK293 binding assays, HEK293T cells were cultured at 37 °C and 5% CO_2_ in DMEM supplemented with 10% FBS and 1% Pen-Strep. Cells were seeded into a six well plate and at ∼50% confluency were transiently transfected using Fugene (Promega) with 500 ng of N-terminal FLAG-tagged MRGPRX2 expression plasmid or empty vector pcDNA negative control, similar to previously described^41^. 48 hours later, cells were removed from the plate with PBS, pelleted, and resuspended in HBS supplemented with 2% FBS and 0.05% bovine serum albumin (BSA). ∼100,000 cells were seeded into a 96-well plate, pelleted, and treated with the indicated concentration of nanobody-Fc for 1 hour or vehicle at 4 °C in duplicate, shaking at 30 rpm. Cells were pelleted and washed with HBS + 2% FBS + 0.05% BSA, then treated with anti-human Fc-AF647 (Biolegend Cat. #410714) for 20 minutes. Cells were pelleted and washed with HBS + 2% FBS + 0.05% BSA, and resuspended in HBS + 2% FBS + 0.05% BSA + 0.4% formaldehyde. Cells were analyzed by cytometry using a CytoFLEX cytometer (Beckman-Coulter), and gated by forward-scatter/side scatter, and singlets (FSC-A versus FSC-H). Mean fluorescence intensity (MFI) at 647 nm within this singlet population was used as the assessment for nanobody binding. Experiments were conducted on two separate days in technical duplicate, and technical duplicates were averaged for each replicate. Nanobodies were tested simultaneously to allow for Bmax and Kd comparisons. Maximal signal was normalized to the 1 μM Nanobody Sim9877-Fc condition. For Kd and Bmax calculations, non-specific signal was MFI at the indicated concentration in HEK293 cells transfected with the empty vector negative control. Data were analyzed in GraphPad Prism version 10.

### β-hexosaminidase release assay

To measure β-hexosaminidase release, ROSA cells were washed twice in warm (37 °C) Tyrode’s buffer (20 mM HEPES with 134 mM NaCl, 5 mM KCl, 1.8 mM CaCl_2_, 1 mM MgCl_2_, 5.5 mM glucose, and 0.3% bovine serum albumin, pH 7.4). Cells were then seeded at 100,000 cells per well (100 μL, 1E6 cells/mL) in a flat bottom clear 96-well plate. Cells were pre-treated with nanobody by adding nanobody at 2x working concentration (100 µM) in Tyrode’s buffer and then cells were incubated for 15 minutes in a humidified incubator with 5% CO_2_ at 37 °C. After the 15 minute incubation, compound 48/80 (Sigma) was added at 50 µM final concentration or nanobody at 50 µM final concentration, with final volumes for all conditions kept at 200 µL. Cells were incubated for 60 minutes in a humidified incubator with 5% CO_2_ at 37 °C. After incubation, samples were transferred to a v-bottom plate, and plates were centrifuged at 500xg for 5 min. For the supernatant-only reading, 50 μL supernatant was added to a new 96-well black flat bottom plate (Nunc) containing 50 μL of 5 μM 4-Methylumbelliferyl-β-D-glucopyranosiduronic acid (4 MUG) (Sigma) diluted in 100 mM citrate buffer (pH 4.5). For the supernatant and lysate reading, cells were lysed by adding 20 μL of 10% Triton-X-100 to each well (0.01% Triton-X-100 final). Samples were mixed vigorously with pipetting to break open the cells. 50 μL of lysate was added to 50 μL of 5 μM 4 MUG. All plates were incubated for 60 min in an incubator without CO_2_ at 37 °C. Lastly, 100 µL of 400 glycine pH 10.7 was added to each well to stop the reaction. Fluorescence intensity at 360/460 was read on a Clariostar (BMG LabTech) with the top optic. Percent degranulation was calculated using the formula: % degranulation= ((supernatant)/(supernatant+lysate))x100.

## REFERENCES

1 Carter, P. J. & Rajpal, A. Designing antibodies as therapeutics. Cell 185, 2789–2805, doi:10.1016/j.cell.2022.05.029 (2022).

2 Banik, S. S. R., Kushnir, N., Doranz, B. J. & Chambers, R. Breaking barriers in antibody discovery: harnessing divergent species for accessing difficult and conserved drug targets. MAbs 15, 2273018, doi:10.1080/19420862.2023.2273018 (2023).

3 Sharma, P., Joshi, R. V., Pritchard, R., Xu, K. & Eicher, M. A. Therapeutic Antibodies in Medicine. Molecules 28, doi:10.3390/molecules28186438 (2023).

4 Morrison, C. Nanobody approval gives domain antibodies a boost. Nat Rev Drug Discov 18, 485–487, doi:10.1038/d41573-019-00104-w (2019).

5 De Pauw, T. et al. Current status and future expectations of nanobodies in oncology trials. Expert Opin Investig Drugs 32, 705–721, doi:10.1080/13543784.2023.2249814 (2023).

6 Lu, R. M. et al. Development of therapeutic antibodies for the treatment of diseases. J Biomed Sci 27, 1, doi:10.1186/s12929-019-0592-z (2020).

7 Harvey, E. P. et al. An in silico method to assess antibody fragment polyreactivity. Nat Commun 13, 7554, doi:10.1038/s41467-022-35276-4 (2022).

8 Mirdita, M. et al. ColabFold: making protein folding accessible to all. Nat Methods 19, 679–682, doi:10.1038/s41592-022-01488-1 (2022).

9 Abramson, J. et al. Accurate structure prediction of biomolecular interactions with AlphaFold 3. Nature, doi:10.1038/s41586-024-07487-w (2024).

10 Jumper, J. et al. Highly accurate protein structure prediction with AlphaFold. Nature 596, 583–589, doi:10.1038/s41586-021-03819-2 (2021).

11 Senior, A. W. et al. Improved protein structure prediction using potentials from deep learning. Nature 577, 706–710, doi:10.1038/s41586-019-1923-7 (2020).

12. 12 Evans, R., et al. Protein complex prediction with AlphaFold-Multimer. bioRxiv 463034 (2022).

13 Watson, J. L. et al. De novo design of protein structure and function with RFdiffusion. Nature 620, 1089–1100, doi:10.1038/s41586-023-06415-8 (2023).

14 Bennett, N. R. et al. Atomically accurate de novo design of single-domain antibodies. bioRxiv, doi:10.1101/2024.03.14.585103 (2024).

15 Skiba, M. A. et al. Antibodies expand the scope of angiotensin receptor pharmacology. Nat Chem Biol, doi:10.1038/s41589-024-01620-6 (2024).

16 Schmid, E. W. & Walter, J. C. Predictomes: A classifier-curated database of AlphaFold-modeled protein-protein interactions. bioRxiv, doi:10.1101/2024.04.09.588596 (2024).

17 Lim, Y. et al. In silico protein interaction screening uncovers DONSON’s role in replication initiation. Science 381, eadi3448, doi:10.1126/science.adi3448 (2023).

18 Basu, S. & Wallner, B. DockQ: A Quality Measure for Protein-Protein Docking Models. PLoS One 11, e0161879, doi:10.1371/journal.pone.0161879 (2016).

19 Lin, Z. et al. Evolutionary-scale prediction of atomic-level protein structure with a language model. Science 379, 1123–1130, doi:10.1126/science.ade2574 (2023).

20 McMahon, C. et al. Yeast surface display platform for rapid discovery of conformationally selective nanobodies. Nat Struct Mol Biol 25, 289–296, doi:10.1038/s41594-018-0028-6 (2018).

21 Cao, C. & Roth, B. L. The structure, function, and pharmacology of MRGPRs. Trends Pharmacol Sci 44, 237–251, doi:10.1016/j.tips.2023.02.002 (2023).

22 Cao, C. et al. Structure, function and pharmacology of human itch GPCRs. Nature 600, 170–175, doi:10.1038/s41586-021-04126-6 (2021).

23 Yang, F. et al. Structure, function and pharmacology of human itch receptor complexes. Nature 600, 164–169, doi:10.1038/s41586-021-04077-y (2021).

24 Staus, D. P. et al. Allosteric nanobodies reveal the dynamic range and diverse mechanisms of G-protein-coupled receptor activation. Nature 535, 448–452, doi:10.1038/nature18636 (2016).

25 McNeil, B. D. et al. Identification of a mast-cell-specific receptor crucial for pseudo-allergic drug reactions. Nature 519, 237–241, doi:10.1038/nature14022 (2015).

26 Azimi, E. et al. Dual action of neurokinin-1 antagonists on Mas-related GPCRs. JCI Insight 1, e89362, doi:10.1172/jci.insight.89362 (2016).

27 Kuehn, H. S., Radinger, M. & Gilfillan, A. M. Measuring mast cell mediator release. Curr Protoc Immunol Chapter 7, Unit7 38, doi:10.1002/0471142735.im0738s91 (2010).

28 Reddy, V. B., Graham, T. A., Azimi, E. & Lerner, E. A. A single amino acid in MRGPRX2 necessary for binding and activation by pruritogens. J Allergy Clin Immunol 140, 1726–1728, doi:10.1016/j.jaci.2017.05.046 (2017).

29 Lansu, K. et al. In silico design of novel probes for the atypical opioid receptor MRGPRX2. Nat Chem Biol 13, 529–536, doi:10.1038/nchembio.2334 (2017).

30 Alkanfari, I., Gupta, K., Jahan, T. & Ali, H. Naturally Occurring Missense MRGPRX2 Variants Display Loss of Function Phenotype for Mast Cell Degranulation in Response to Substance P, Hemokinin-1, Human beta-Defensin-3, and Icatibant. J Immunol 201, 343–349, doi:10.4049/jimmunol.1701793 (2018).

31 Tatemoto, K. et al. Immunoglobulin E-independent activation of mast cell is mediated by Mrg receptors. Biochem Biophys Res Commun 349, 1322–1328, doi:10.1016/j.bbrc.2006.08.177 (2006).

32 Al Hamwi, G. et al. MAS-related G protein-coupled receptors X (MRGPRX): Orphan GPCRs with potential as targets for future drugs. Pharmacol Ther 238, 108259, doi:10.1016/j.pharmthera.2022.108259 (2022).

33 Shtessel, M. et al. MRGPRX2 Activation Causes Increased Skin Reactivity in Patients with Chronic Spontaneous Urticaria. J Invest Dermatol 141, 678–681 e672, doi:10.1016/j.jid.2020.06.030 (2021).

34 Hauser, A. S., Attwood, M. M., Rask-Andersen, M., Schioth, H. B. & Gloriam, D. E. Trends in GPCR drug discovery: new agents, targets and indications. Nat Rev Drug Discov 16, 829–842, doi:10.1038/nrd.2017.178 (2017).

35 Yang, D. et al. G protein-coupled receptors: structure-and function-based drug discovery. Signal Transduct Target Ther 6, 7, doi:10.1038/s41392-020-00435-w (2021).

36 Graff, D. E., Shakhnovich, E. I. & Coley, C. W. Accelerating high-throughput virtual screening through molecular pool-based active learning. Chem Sci 12, 7866–7881, doi:10.1039/d0sc06805e (2021).

37 Dunbar, J. & Deane, C. M. ANARCI: antigen receptor numbering and receptor classification. Bioinformatics 32, 298–300, doi:10.1093/bioinformatics/btv552 (2016).

38 Skiba, M. A. et al. Antibodies Expand the Scope of Angiotensin Receptor Pharmacology. bioRxiv 23.554128 (2023).

39 Martin, W. L., West, A. P., Jr., Gan, L. & Bjorkman, P. J. Crystal structure at 2.8 A of an FcRn/heterodimeric Fc complex: mechanism of pH-dependent binding. Mol Cell 7, 867–877, doi:10.1016/s1097-2765(01)00230-1 (2001).

40 Smith, J. S. et al. The M3 Muscarinic Acetylcholine Receptor Can Signal through Multiple G Protein Families. Mol Pharmacol 105, 386–394, doi:10.1124/molpharm.123.000818 (2024).

41 Eiger, D. S. et al. Phosphorylation barcodes direct biased chemokine signaling at CXCR3. Cell Chem Biol 30, 362–382 e368, doi:10.1016/j.chembiol.2023.03.006 (2023).

42 Olsen, R. H. J. et al. TRUPATH, an open-source biosensor platform for interrogating the GPCR transducerome. Nat Chem Biol 16, 841–849, doi:10.1038/s41589-020-0535-8 (2020).

43 Saleh, R. et al. A new human mast cell line expressing a functional IgE receptor converts to tumorigenic growth by KIT D816V transfection. Blood 124, 111–120, doi:10.1182/blood-2013-10-534685 (2014).

