## Supplementary Information for "*In silico* discovery of nanobody binders to a G-protein coupled receptor using AlphaFold-Multimer"

**Supplementary Table 1. Nanobody Sequences.** CDRs are shown in red.

| **Name** | **Sequence**<br>(Bold = mutation from corresponding parent, red italics = CDR regions) |
|---|---|
| Sim8619 (MRGPRX2 Rank 1) | QVQLQESGGGLVQAGGSLRLSCAASGSIFYIRGMGWYRQAPGKERELVAGIDVGAITYYADSVKGRFTISRDNAKNTVYLQMNSLKPEDTAVYYCAVWAYTRAGYTTVYAYWGQGTQVTVSS |
| Sim8619 R102A | QVQLQESGGGLVQAGGSLRLSCAASGSIFYIRGMGWYRQAPGKERELVAGIDVGAITYYADSVKGRFTISRDNAKNTVYLQMNSLKPEDTAVYYCAVWAYT**A**AGYTTVYAYWGQGTQVTVSS |
| Sim7252 (MRGPRX2 Rank 2) | QVQLQESGGGLVQAGGSLRLSCAASGSISRWLGMGWYRQAPGKEREFVAGITSGANTNYADSVKGRFTISRDNAKNTVYLQMNSLKPEDTAVYYCAAHYVVSVILVYWGQGTQVTVSS |
| Sim0563 (MRGPRX2 Rank 3) | QVQLQESGGGLVQAGGSLRLSCAASGTIFLPSSMGWYRQAPGKERELVAGITYGAITYYADSVKGRFTISRDNAKNTVYLQMNSLKPEDTAVYYCAVVGLGYGWHFYWGQGTQVTVSS |
| Sim9877 (MRGPRX2 Rank 5) | QVQLQESGGGLVQAGGSLRLSCAASGYISSFPVMGWYRQAPGKERELVAAIGSGGITYYADSVKGRFTISRDNAKNTVYLQMNSLKPEDTAVYYCAVAGYNIGSYYYWGQGTQVTVSS |
| Sim4717 (MRGPRX2 Rank 6) | QVQLQESGGGLVQAGGSLRLSCAASGNIFFYPDMGWYRQAPGKEREFVATIGGGGITYYADSVKGRFTISRDNAKNTVYLQMNSLKPEDTAVYYCAVGGIYVGPHVYWGQGTQVTVSS |
| Sim4784 (MRGPRX2 Rank 7) | QVQLQESGGGLVQAGGSLRLSCAASGTISPPTYMGWYRQAPGKERELVASIGAGSNTNYADSVKGRFTISRDNAKNTVYLQMNSLKPEDTAVYYCAAIFGRLWYHLYWGQGTQVTVSS |
| Sim3014 (MRGPRX2 Rank 31) | QVQLQESGGGLVQAGGSLRLSCAASGYISRLGLMGWYRQAPGKEREFVAAISLGSTTYYADSVKGRFTISRDNAKNTVYLQMNSLKPEDTAVYYCAVYNQRRIVDYSNFAYWGQGTQVTVSS |
| Sim4177 (MRGPRX2 Rank 90) | QVQLQESGGGLVQAGGSLRLSCAASGSISRRGWMGWYRQAPGKEREFVATITFGASTYYADSVKGRFTISRDNAKNTVYLQMNSLKPEDTAVYYCAVFYSQYWLYLYWGQGTQVTVSS |
| Sim1846 (MRGPRX2 Rank 121) | QVQLQESGGGLVQAGGSLRLSCAASGYIFNATVMGWYRQAPGKERELVATITGGTNTYYADSVKGRFTISRDNAKNTVYLQMNSLKPEDTAVYYCAVVTFVVIPYTYWGQGTQVTVSS |
| Sim7492 (MRGPRX2 Rank 151) | QVQLQESGGGLVQAGGSLRLSCAASGNIFRIGPMGWYRQAPGKERELVATIASGAITYYADSVKGRFTISRDNAKNTVYLQMNSLKPEDTAVYYCAANHWVASFWYRYLEYWGQGTQVTVSS |
| Nanobody 60 (Negative Control) | QVQLQESGGGLVQAGGSLRLSCAASGSIFSLNDMGWYRQAPGKLRELVAAITSGGSTKYADSVKGRFTISRDNAKNTVYLQMNSLKAEDTAVYYCNAKVAGTFSIYDYWGQGTQVTVSS |

### SUPPLEMENTARY FIGURES

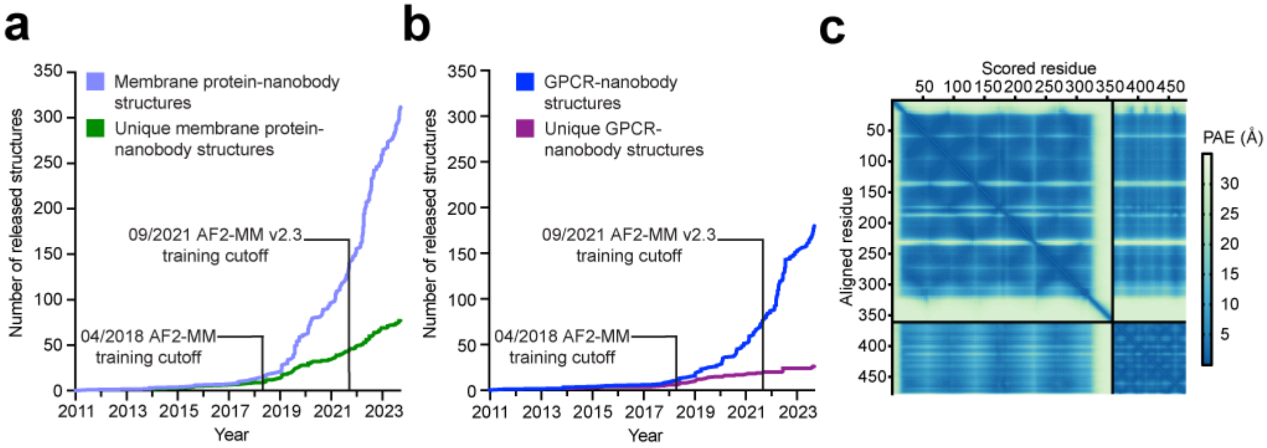


**Supplementary Figure 1.** a-b). The number of membrane protein-nanobody structures and GPCR-nanobody structures deposited in the PDB over time. c). PAE plot of the AT1R/AT118 complex AF-M prediction. The low PAE values in the AT1R/AT118 binding interface suggest AF-M is highly confident in the binding interaction.

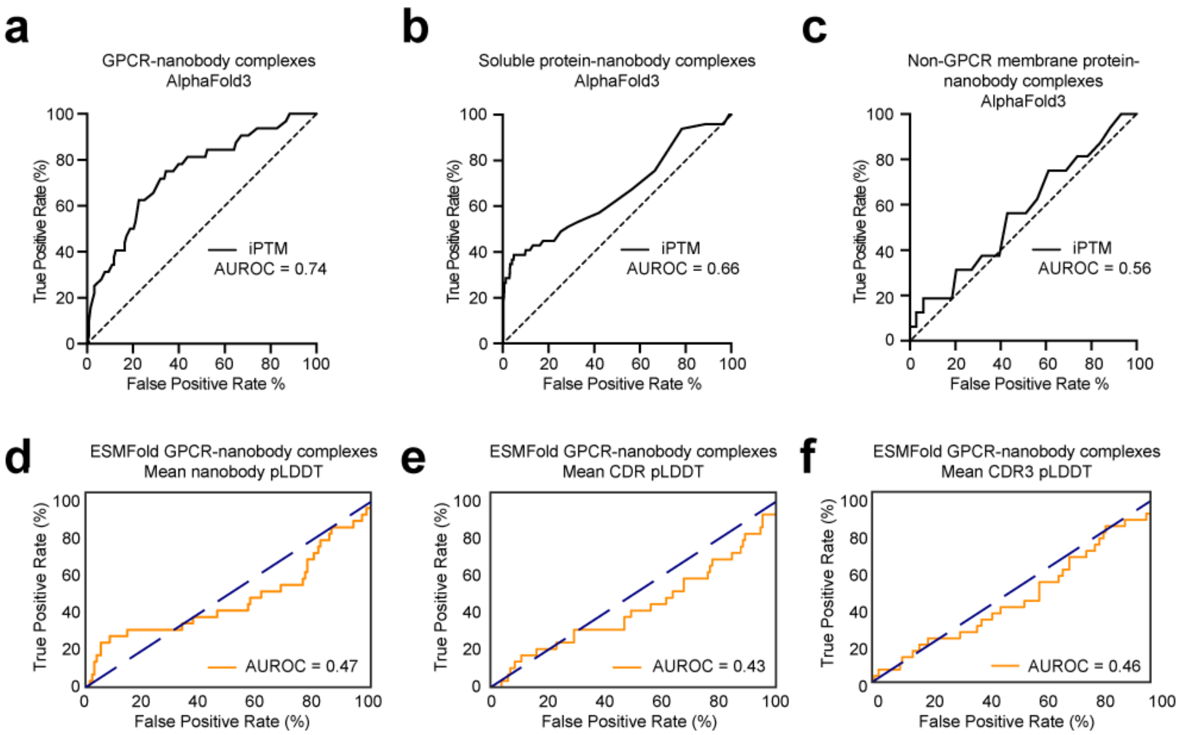


**Supplementary Figure 2.** (a-c). AlphaFold3 can differentiate between true GPCR-nanobody complexes versus negative controls (a) and more moderately, true soluble protein-nanobody complexes and negative controls (b). AlphaFold3 cannot differentiate between real non-GPCR membrane protein-nanobody complexes and negative controls (c). d-f). ESMFold does not accurately differentiate between true GPCR-nanobody binding interactions and negative control interactions, as assessed by pLDDT values of the entire nanobody (d), the CDR regions (e), and the CDR3 region (f).

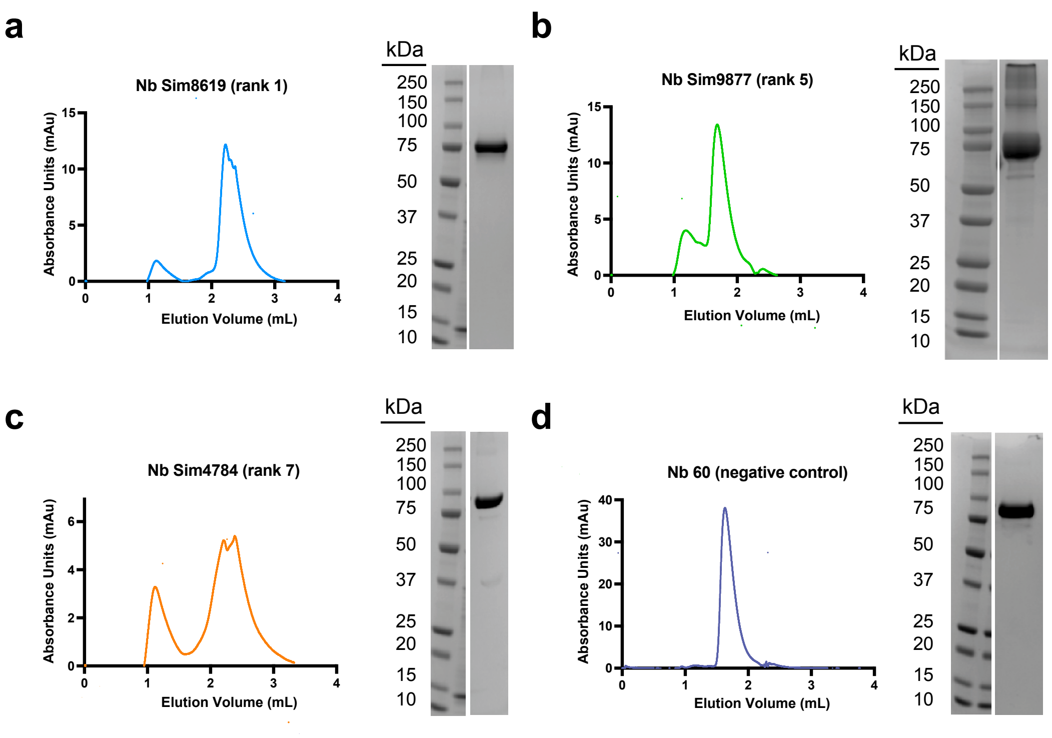


**Supplementary Figure 3.** a-d). Top-ranked MRGPRX2 simulated nanobodies and the negative control nanobody 60 were expressed and purified from Expi293 cells and then analyzed by analytical size exclusion chromatography, gel electrophoresis, and Coomassie blue staining to assess monodispersity and purity. Nanobodies were run on a Superdex 200 Increase 3.2/300 column.

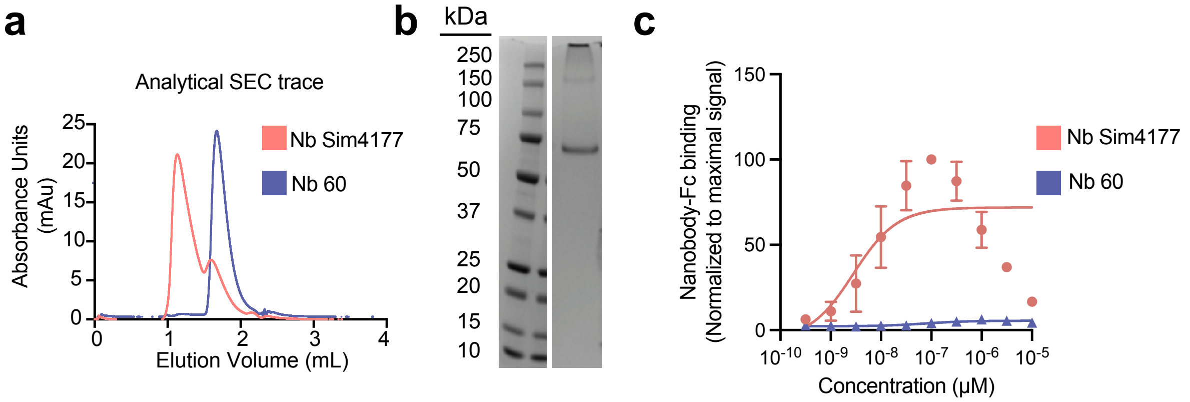


**Supplementary Figure 4.** a) Size exclusion chromatography trace of nanobody Sim4177 (rank 90) overlayed with the trace of nanobody 60, showing that most of Sim4177 elutes near the void of the column. Nanobodies were run on a Superdex 200 Increase 3.2/300 column. b) Coomassie stained gel of purified nanobody Sim4177, showing the presence of a higher molecular weight species. c) Nanobody Sim4177 binds to ROSA mast cells, but at high concentrations, exhibits lower binding, potentially indicating aggregation. In Panel C, experiments were performed in biological duplicate with error bars representing mean +/- SEM for technical replicates of a representative experiment.

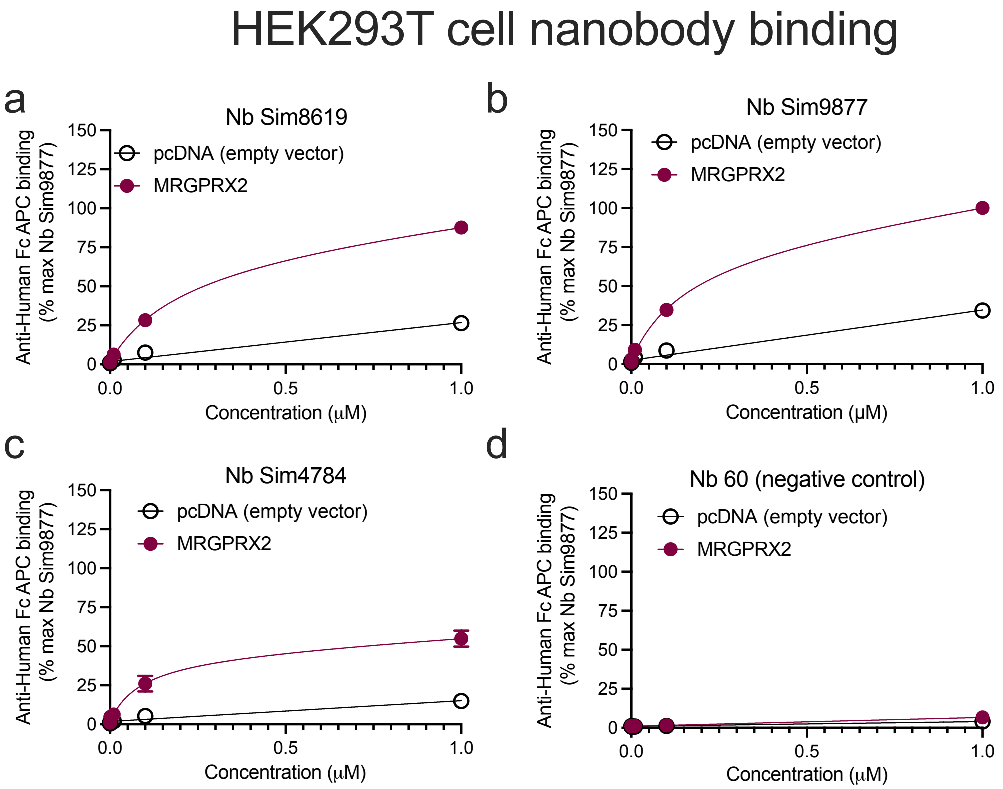


**Supplemental Figure 5**. HEK293T cells transiently transfected with either human MRGPRX2 or empty vector (pcDNA) were treated for 1 hour at the indicated concentration of the Fc-conjugated a) Sim8619 (rank 1), b) Sim9877 (rank 5), c) Sim4784 (rank 7) or d) the negative control nanobody 60. Data were normalized to maximal mean fluorescence intensity signal. Experiments were conducted in duplicate on separate days, with two technical replicates merged per replicate. Dissociation constant and max binding data are available in Table 1 of the main text.

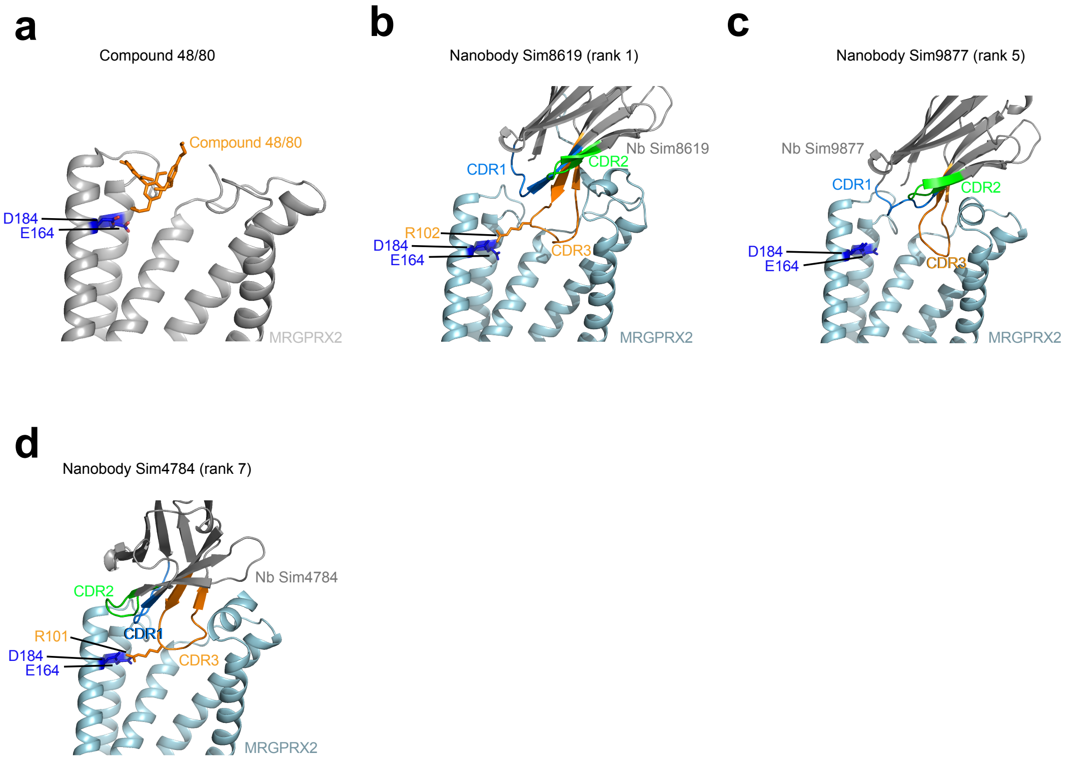


**Supplementary Figure 6.** a) Structure of Compound 48/80 bound to MRPGRX2 (PDB 7VV6) showing that a positively charged side group of Compound 48/80 interacts with two acidic MRGPRX2 residues. b-d) AlphaFold2 predictions of candidate simulation nanobodies bound to MRGPRX2. Nanobody Sim8619 (rank 1) and Nanobody 4784 (rank 7) both possess arginine residues in their CDR3 domains that interact with the same two acidic residues in MRGPRX2 that Compound 48/80 interacts with.

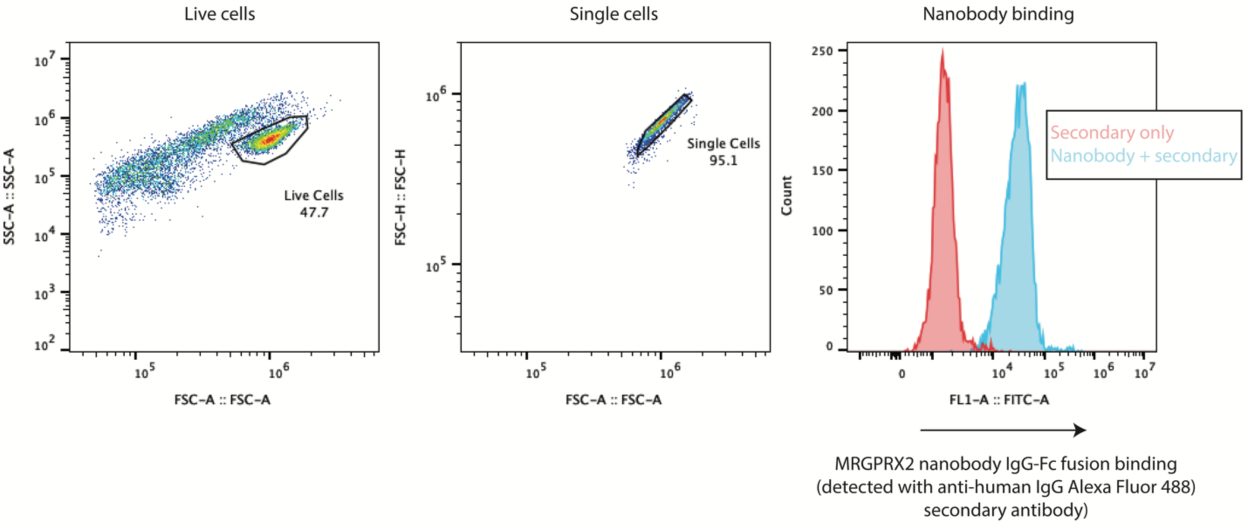


**Supplementary Figure 7.** Representative gating strategy for mammalian cells. Cells were first gated on live cells and then on singlet cells before analysis of nanobody binding.
